# Insights from *Bacillus anthracis* strains isolated from permafrost in the tundra zone of Russia

**DOI:** 10.1101/486290

**Authors:** V Timofeev, I Bahtejeva, R Mironova, G Titareva, I Lev, D Christiany, A Borzilov, A Bogun, G Vergnaud

**Affiliations:** State Research Center for Applied Microbiology & Biotechnology» (FBIS SRCAMB), Russia; Institute for Integrative Biology of the Cell (I2BC), CEA, CNRS, Univ. Paris-Sud, Université Paris-Saclay, 91198, Gif-sur-Yvette cedex, France

**Keywords:** *Bacillus anthracis*, tundra region, MLVA-genotyping, whole genome SNP analysis

## Abstract

This article describes *Bacillus anthracis* strains isolated during an outbreak of anthrax on the Yamal Peninsula in the summer of 2016 and independently in Yakutia in 2015. A common feature of these strains is their conservation in permafrost, from which they were extracted either due to the thawing of permafrost (Yamal strains) or as the result of paleontological excavations (Yakut strains). All strains isolated on the Yamal share an identical genotype belonging to lineage B.Br.001/002, pointing to a common source of infection in a territory over 250 km in length. In contrast, during the excavations in Yakutia, three genetically different strains were recovered from a single pit. One strain belongs to B.Br.001/002, as the Yamal strains. Despite the remoteness of Yamal from Yakutia, whole genome sequence analysis showed that the B.Br.001/002 strains are very closely related. The two other strains contribute to two different branches of A.Br.008/011, one of the remarkable polytomies described so far in *B. anthracis* population. The geographic distribution of the strains belonging to this polytomy is suggesting that this polytomy emerged in the thirteenth century, in combination with the constitution of a unified Mongol empire extending from China to Eastern Europe. We propose an evolutionary model for *B. anthracis* recent evolution in which the B lineage spread throughout Eurasia and was subsequently replaced by the A lineage except in some geographically isolated areas.

## Introduction

The etiological agent of the anthrax disease is the gram-positive bacterium *Bacillus anthracis*. A key feature of this microorganism, which largely determines its epidemiological potential and population structure, is the ability to form endospores, extremely resistant to adverse environmental factors and able to remain viable for a long time [1–12]. High preservation of spores explains that even in regions where this disease has not been observed for decades, disease outbreaks are possible, leading to significant economic damage, the mass mortality of livestock, and human victims. In addition, due to the high virulence of *B. anthracis*, the stability of endospores in the environment and the simplicity of cultivation, this bacterium is considered a potential biological weapon or tool for bioterrorism [13, 14], as illustrated by the anthrax contaminated letters in 2001 in the USA [15].

Anthrax is now very rare in most European countries [16] but remains a significant problem mainly in sub-Saharan Africa and in some regions of Asia [17–20]. Anthrax is endemic in Russia, where the disease manifests itself as sporadic cases among animals and rare cases of the disease among the population [21]. The presence of large territories hosting populations of wild and domestic ungulates creates a favorable context for disease outbreaks of epizootics, and the low human population density in most parts of the country makes it difficult to conduct anti-epidemiс measures, and to correctly account for anthrax animal burial sites. Past animal burial sites are often not documented and occasionally corpses were not buried. These burial grounds, and entire territories, where previously epizootics took place, may become involved in increased economic activities, which, given the potentially high preservation of *B. anthracis* spores in a cold environment, could lead to new outbreaks of the disease [22]. Of particular interest in this regard is the tundra zone of Russia, located between 55 and 68 degrees north latitude.

One of the features of this climatic zone is the presence of permafrost. Permafrost is defined as lithosphere material (soil and sediment) permanently exposed to temperatures ≤0 °C and remaining frozen for at least two consecutive years. Permafrost can extend down to more than 1000 m in depth and remain frozen for thousands of years [23, 24]. In permafrost conditions, the preservation of microorganisms can significantly increase, and thus permafrost is a peculiar accumulator of microbiota [25, 26]. The preservation of spores of bacilli, including *B. anthracis*, in the permafrost theoretically should significantly exceed the preservation of microorganisms in the vegetative form. Consequently, permafrost might allow the discovery of archaic forms of this microorganism, which could supplement our knowledge of the evolution of the anthrax microbe. We investigated strains of *B. anthracis* isolated in two tundra zones - during the outbreak of anthrax in Yamal in the summer of 2016 and during extraction of paleontological material from the permafrost in Yakutia in 2015.

## Materials and Methods

### Yamal samples

In the summer of 2016 an outbreak of anthrax occurred on the Yamal Peninsula. The previous outbreak of anthrax was registered in 1941 and the district was officially declared “anthrax-free” territory of the USSR since 1968. In 2007, the compulsory vaccination of reindeer was abandoned. On July 16^th^ 2016, the United Duty Control Service of the Yamal District was informed of deer death by private reindeer herders. The deer’s deaths began at the estuary of the Nerosaveyakha River near Lake Pisyoto. Reindeer herders reported that sick animals became sluggish, began to move slowly, then fell and quickly died. No ulcers or skin lesions could be detected. On July 17^th^ and 18^th^, the Veterinary Service of the Yamal-Nenets Autonomous Area arrived in the area for clinical examination of animals and autopsy. Pathological material was sent to the Tyumen Regional Veterinary Laboratory. An autopsy showed cardiac and pulmonary insufficiency and the preliminary diagnosis was death from a heat stroke as July 2016 had anomalously hot weather, with temperatures above 35°C. Complementary investigations by veterinarians sent to reindeer herders camps on July 19-29 led the Tyumen Regional Veterinary Laboratory to report a suspicion of *B. anthracis* on July 24^th^ and to take prophylactic measures (vaccination and chemotherapy, restrictions on animal movements). Additional samples were sent to the All-Russian Scientific Research Institute of Veterinary Virology and Microbiology (ARSRIVVM) in Pokrov (Moscow region).

By this time, the disease was observed in three focuses: Lake Pisyoto area, Novoportovskaya tundra, Evayakha River area. The outbreak sites were separated by distances up to 250 km, including two water barriers - the Gulf of Ob (width from 30 to 80 km) and the Taz Estuary (average width is about 25 kilometers).

On July 25^th^, a complete laboratory confirmation of presence of *B. anthracis* in samples taken from dead deer was obtained by ARSRIVVM. A pure culture of *B. anthracis* was isolated from one sample, this strain was called 5875. The Governor of Yamal-Nenets Autonomous District introduced a quarantine regime in the Yamal district. SRCAMB’s and ARSRIVVM’s employees went to Salekhard to sample and to consult local sanitary and medical institutions. Specialists of the Stavropol Anti-Plague Institute arrived on July 26th.

Starting on July 26, arrived experts organized a diagnostic laboratory in the Center for Hygiene and Epidemiology in the Yamal-Nenets Autonomous District. During the outbreak, samples from people potentially infected were investigated by this diagnostic laboratory. Medical authorities decided to hospitalize to Salekhard all children from the outbreak areas even without visible signs of the disease. People evacuation to temporary camps equipped by that time were started and preventive antibiotic therapy was applied. SRCAMB’s experts flew from Salekhard to the disease focus in the Lake Pisyoto area to survey the local population and to collect samples. Until this time, both the local population and veterinarians working in this outbreak area were skeptical about the possibility of anthrax, and favored the heat shock hypothesis as a number of other infections harmless for humans could have caused the death of animals weakened by heat. Furthermore simultaneously with the beginning of vaccination and antibiotic therapy, the temperature of the infection focus decreased sharply, so there were reasons to believe that the cessation of new cases of the disease resulted from the lowering of temperature.

The typical development of the disease was as follows: a seemingly healthy deer would become suddenly weak, unable to walk and forced to lay down a few hours later, and would die after a few additional hours. In most cases the nose was bleeding (sometimes the anus too), rigor mortis developed at the usual time.

SRCAMB’s experts took samples of soil, water, blood samples, ears, and lymph nodes of dead deer. The samples were delivered to Salekhard on July 27^th^, and eventually to SRCAMB on July 28th. On July 29^th^, strain 5875 was sent from ARSRIVVM to SRCAMB. On the same day, SRCAMB received a strain isolated from a sick person (washed off from skin infection) subsequently called Yamal_12 and insects caught by veterinarians working in the outbreak area: nine *Scopeuma stercorarium* and four *Hydrotaca dentipes* from Salekhard. On August 13^th^, SRCAMB received strain 6063 isolated in epidemic area by ARSRIVVM.

### Yakutia samples

On August 12, 2015, miners extracting mammoth tusks from the permafrost on the bank of the river Uyandina in the Abyisk ulus (district) of Yakutia 57 km from the district center Belaya Gora (“White Mountain”) (latitude N (68.564567), longitude E (144.769827)) found two kittens of the cave lion *Panthera leo spelaea* frozen in ice. The discovery was remarkable by its unprecedented degree of preservation - the animals preserved wool and soft tissues. The bodies of the kittens were transferred to paleontologists. Some samples were taken for microbiological examination to Institute of Oil and Gas of the Siberian Branch of the Russian Academy of Sciences in Yakutsk (IPMR SB RAS), the nearest scientific institution. June 01, 2016 an unknown bacillus-like strain was isolated in the laboratory of geochemistry of caustobioliths of IPMR SB RAS and sent to the Institute of Genetics and Selections of Industrial Microorganisms (GosNIIgenetika) in Moscow for identification. September, 01 this strain was identified as *B. anthracis* according to PCR results. Since this institute is not equipped to carry out work with pathogenic microorganisms and has no appropriate license, this initial culture was destroyed. September, 21 according to order of the Chief State Sanitary Doctor of Yakutia, soil samples were collected in paleontological discovery point. Six separately packed glass jars with soil samples (200 g each), taken from a depth of 1 to 6 meters with an interval of 1 meter were received by SRCAMB in December 2016.

## Animal experiments

### Ethics statement

All protocols for animal experiments were approved by the State Research Center for Applied Microbiology and Biotechnology Bioethics Committee (Permits No: VP-2016/4 and VP-2016/5). They were performed in compliance with the NIH Animal Welfare Insurance #A5476-01 issued on 02/07/2007 and the European Union guidelines and regulations on handling, care, and protection of laboratory animals (http://ec.europa.eu/environment/chemicals/lab_animals/home_en.htm).

All used animals were purchased from Laboratory Animals Breeding Center, Shemyakin and Ovchinnikov Institute of Bioorganic Chemistry, Russia. They were housed in polycarbonate cages with space for comfortable movement (5 mice per 484 cm^2^ cage or 2-3 guinea pigs per 864 cm^2^ cage) and easy access to food and water, under constant temperature (22°C ± 2°C) and humidity conditions (50% ± 10%) and a 12-hour light/12-hour dark cycle.

Approved protocols provided scientifically validated humane endpoints, including pre-set criteria for euthanasia of moribund animals by CO_2_ inhalation. Animals were euthanized when they became lethargic, dehydrated, moribund, unable to rise, or non-responsive to touch. Surviving animals were euthanized after the observation period.

### Mice

Six-to-eight-weeks-old BALB/C mice of both genders, weighing 18–20g were used in all our experiments. They were fed Mouse Mixed Fodder PK-120 (Laboratorkorm, Russia) and provided tap water *ad libitum* throughout observation period.

For virulence evaluation mice were randomly divided in four groups of ten animals and infected s.c. by doses:10^0^, 10^1^, 10^2^ and 10^3^ spores/animal. They were observed for 30 days after infection.

For bioassay we used groups of three mice for each tested sample. Mice were inoculated subcutaneously with sample dispended in 0.3 ml of PBS in the inner part of the upper thigh. Animals were observed during 10 days, after which surviving mice were euthanized. Dead and euthanized mice were necropsied, and samples of their spleens and livers were inoculated on Petri dishes with selective anthrax agar (SRCAMB).

### Guinea pigs

Guinea pigs were used for evaluating the virulence of strains isolated in Yamal epidemic area. We used five-to-seven-weeks-old animals of both genders, weighing 350–450g. They were fed Granuled Fodder KK-122 (Laboratorkorm, Russia) and provided tap water *ad libitum* during the entire experiment (30 days). Guinea pigs were randomly divided in three groups of five animals and infected s.c. by doses of 10^2^, 10^3^, 10^4^ spores/animal.

### Bacterial culture and DNA extraction

For bacterial cultures isolation, cultivation for DNA extraction, and for lecitinase and hemolytic activity evaluation we used GRM agar, selective anthrax agar, yolk agar and blood agar, manufactured in SRCAMB, Russia.

DNA from field and clinical samples was isolated using «Reagent kit «K-Sorb» for DNA extraction on microcolumns» (Syntol, Russia). DNA from bacterial cultures was isolated using GenElute™ Bacterial Genomic DNA Kit (Sigma-Aldrich, USA).

### PCR analyses

PCR amplifications were run on the CFX96 ™ Real-Time PCR Detection System (Bio-Rad Laboratories, Inc, USA). For MLVA and canSNP genotyping we used 2.5× PCRmix M-427 with SYBR-GreenI (Syntol, Moscow, Russia). Primers were synthesized by Syntol, Russia.

PCR detection of *B. anthracis* DNA was performed using Real-Time PCR-test systems «MULTI-FLU» (SRCAMB, Obolensk, Russia) and «OM-screen-anthrax-RT» (Syntol, Moscow, Russia).

MLVA was performed using primers as indicated in Thierry et al. [27], but using monoplex PCR. PCR products size was evaluated using agarose gel-electrophoresis. The PCR products and a 20 bp ladder (Bio-Rad, USA) were electrophoresed at 100 V for 240 min on a 32-cm length 3% agarose gel prepared in 0.5× TBE. The DNA fragments were visualized with ethidium bromide staining and ultraviolet (254 nm) using the Doc-Print gel documenting system and PhotoCaptMw software version 99.04 (Vilber Lourmat, Marne-la-Vallée, France). PCR products larger than 600 bp were reanalyzed on 2% agarose gel for better resolution. Also in these few cases we confirmed the size of amplicon using Experion™ Automated Electrophoresis System (BioRad, Hercules, USA) and by sequencing the fragment.

canSNP-genotyping was performed as described in [28].

### Whole Genome SNP Analysis

Yamal strains DNA was sequenced using the Ion Torrent PGM (Life Technologies, USA). Ion PGM Reagents 400 Kit (Life Technologies, USA) and Ion 318 Chip Kit (Life Technologies, USA) were used for sequencing. For each genome, reads were de novo assembled using 2.9 Newbler assembler (Roche).

Yakutia strains whole-genome sequencing was performed using the Illumina MiSeq instrument. DNA libraries were prepared using Nextera DNA Library Preparation Kit. Miseq Reagent Kit v3 was used for sequencing. For each genome, reads were assembled de novo using SPAdes v. 3.9 (http://bioinf.spbau.ru/spades).

Additional sequence read archives were recovered from the European Nucleotide Archive (ENA) using the enaBrowserTools (https://github.com/enasequence/enaBrowserTools) and the Aspera (Aspera, Inc., USA) high-speed file transfer protocol. Genome assemblies were downloaded from NCBI and *in silico* converted into 100 bp. sequence reads files with a 50x coverage using a homemade python script. Sequencing reads were mapped on reference genome *B. anthracis* Ames ancestor assembly GCA_000008445 using BioNumerics version 7.6.3. SNPs were called within BioNumerics using the strict closed dataset option. Minimum spanning trees were produced allowing the creation of hypothetical missing links.

## Results

### Investigation of the Yamal samples

PCR-diagnostic of soil, water and necropsy samples showed that all samples (n=23) except soil from the reindeer herding camp (n=5) contained DNA of *B. anthracis*. Set of nutrient medias - GRM agar, selective anthrax medium, yolk agar, blood agar was inoculated with materials from investigated samples. Typical, *B. anthracis*-like colonies grew from all PCR positive samples, none from PCR-negative samples. Microscopic studies of these colonies showed the presence of chains of gram-positive bacillus coated with a capsule. All these bacillus-like colonies were PCR positive for *B. anthracis*. Some colonies of extraneous microflora grew from these samples. All insects samples were PCR-negative, and no colonies of *B. anthracis* or extraneous microflora could be recovered. This negative result is likely due to the use of ethanol for better preservation of entomological specimens. Consequently, we were unable to confirm or disprove the potential role of bloodsucking insects in the spreading of the disease over long distances.

Soil from the place of death, soil from the camp, water from a nearby lake, cervical lymph node, blood from the neck region, blood discharge from the anus, and all the clinical samples were used in a bioassay. All the mice, (except mice infected by soil from the camp) died with symptoms of anthrax on the second day after infection, their spleen and injection sites contained live anthrax bacteria as shown by inoculation on Petri dishes.

MLVA genotyping was applied in order to establish whether several genotypes circulated in the epidemic zone. We initially used loci vrrA, Bams03, Bams05, Bams22, Bams44, and vntr23 derived from the MLVA7 scheme proposed by Thierry et al. [27]. This set of loci allowed to genotype all isolates during one day with a high degree of reliability and proved to be very useful in conducting an epidemiological investigation, when it is required to minimize the time spent on analysis, down to hours. Later it turned out that the same MLVA profile was found in all strains isolated in the summer of 2016 on the Yamal, regardless of the isolation place (Lake Pisyoto area, Novoportovskaya tundra, Evayakha River area) and of the institution by which the samples were analyzed (S1 table) even when using 25 loci. Querying the *Bacillus anthracis* v4_0 MLVA database at http://microbesgenotyping.i2bc.paris-saclay.fr [27] indicates that the Yamal strains belong to the B-clade [29]. All genotypes present in the MLVA database differed at four loci or more among the 25 loci. canSNP-genotyping using melt-curves [28] assigned all Yamal strains to B.Br.001/002 lineage in agreement with the MLVA genotype.

The finding of a unique MLVA25 profile suggested that one strain circulated throughout the epidemic, possibly from a unique source of infection of humans and animals. No strains in collections of SRCAMB and Stavropol Anti-Plague Institute (Reference Center for the control of Anthrax) showed the same MLVA25 profile.

We selected four strains collected in Lake Pisyoto outbreak area for further work including whole genome sequencing (table 1).

**Table 1.**
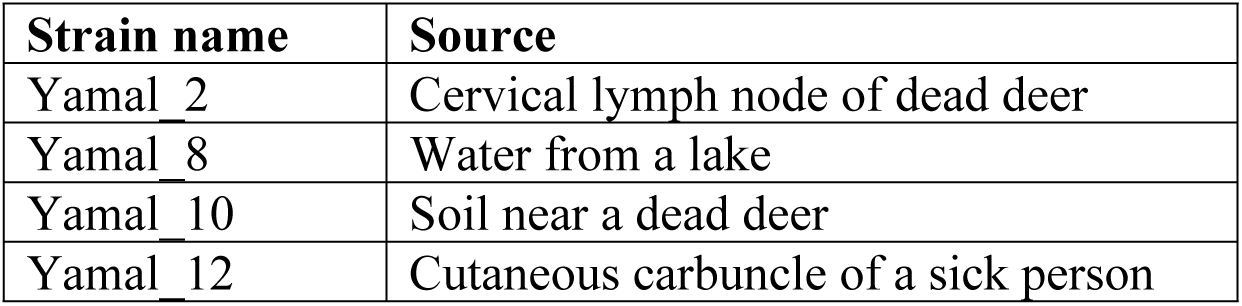
*B. anthracis* strains isolated from Yamal outbreak field and clinical samples

The strains listed in table 1 were typical for a combination of phenotypic properties, but the clinical isolate Yamal_12 initially had lecithinase activity, unlike the other isolates. After two passages on solid media, this activity was lost.

### Investigation of the Yakoutia samples

Several typical *B. anthracis* colonies were cultivated from soil samples from a depth of 2, 3 and 4 meters. The colonies were confirmed as *B. anthracis* by MLVA analysis at seventeen loci (MLVA17). Three genotypes, subsequently called 3Ya, 4Ya, 5Ya could be distinguished (see S1 table). Genotype 4Ya was equally represented in samples from a depth of 2 and 3 m (Table 2). Genotypes 3Ya and 4Ya differ at four loci but querying the *B. anthracis* MLVA database indicates that both are closest to strains belonging to canSNP clade A.Br.008/009. MLVA17 genotype 5Ya is identical to the Yamal MLVA genotype.

**Table 2.**
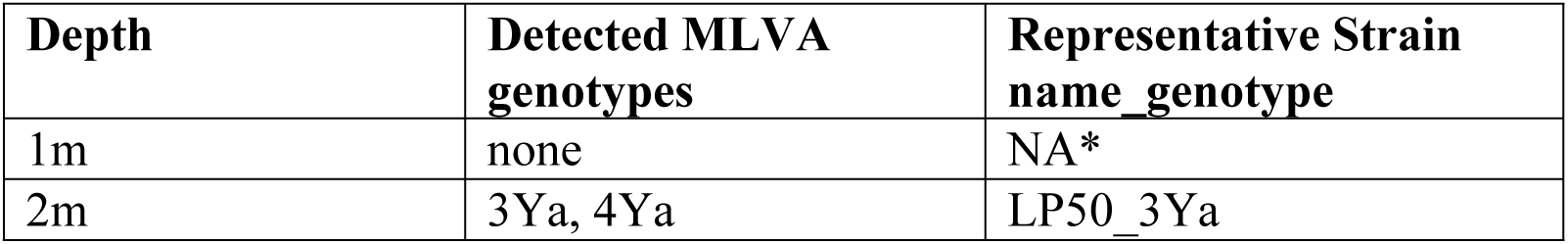
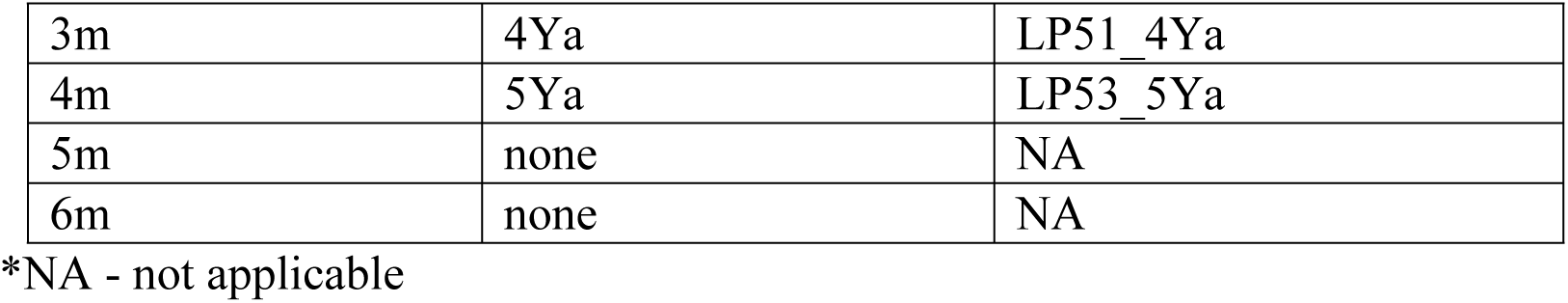
Yakutia soil samples investigations and strains selected for whole genome sequencing

### Whole genome SNP analysis

Contrary to our expectations, the recovered MLVA genotypes are very similar to already know genotypes. Furthermore, the Yamal MLVA genotype is identical to one of the three Yakut strains genotypes. Finally, we recovered three different MLVA genotypes from the same spot in Yakutia, whereas the Yamal 2016 outbreak was associated with a unique genotype in spite of its wide geographic dispersion. One simple explanation for this surprising observation from the Yakut excavation could be contamination when analyzing the samples in SRCAMB. The resistance of the endospores is a well-known cause of accidental laboratory contaminations as recently recalled [30]. In order to investigate this possibility we sequenced four Yamal (Table 1) and three Yakut (Table 2) strains, together with the four strains in the SRCAMB collection showing the closest MLVA genotype. We performed a whole genome SNP analysis comparison of these ten genomes and also included archive reads and assemblies downloaded from EBI-ENA. We used *B. cereus* strain ISSFR-23F representative of one *B. cereus* lineage closest to *B. anthracis* as outgroup to root the tree [31]. Figure 1 shows the relative position of the Yamal and Yakut strains within the global *B. anthracis* population represented by a selection of 50 strains among 650. The longest genetic distance links the MRCA of the *B. anthracis* species and the *B. cereus* outgroup. The position of the ancestor of the *B. anthracis* species along this branch is unknown. The four Yamal strains are identical and belong to B.Br001/002 in agreement with the canSNP typing. The Yakut Ya5 strain is the closest neighbor but is clearly distinct.

**Fig 1.**
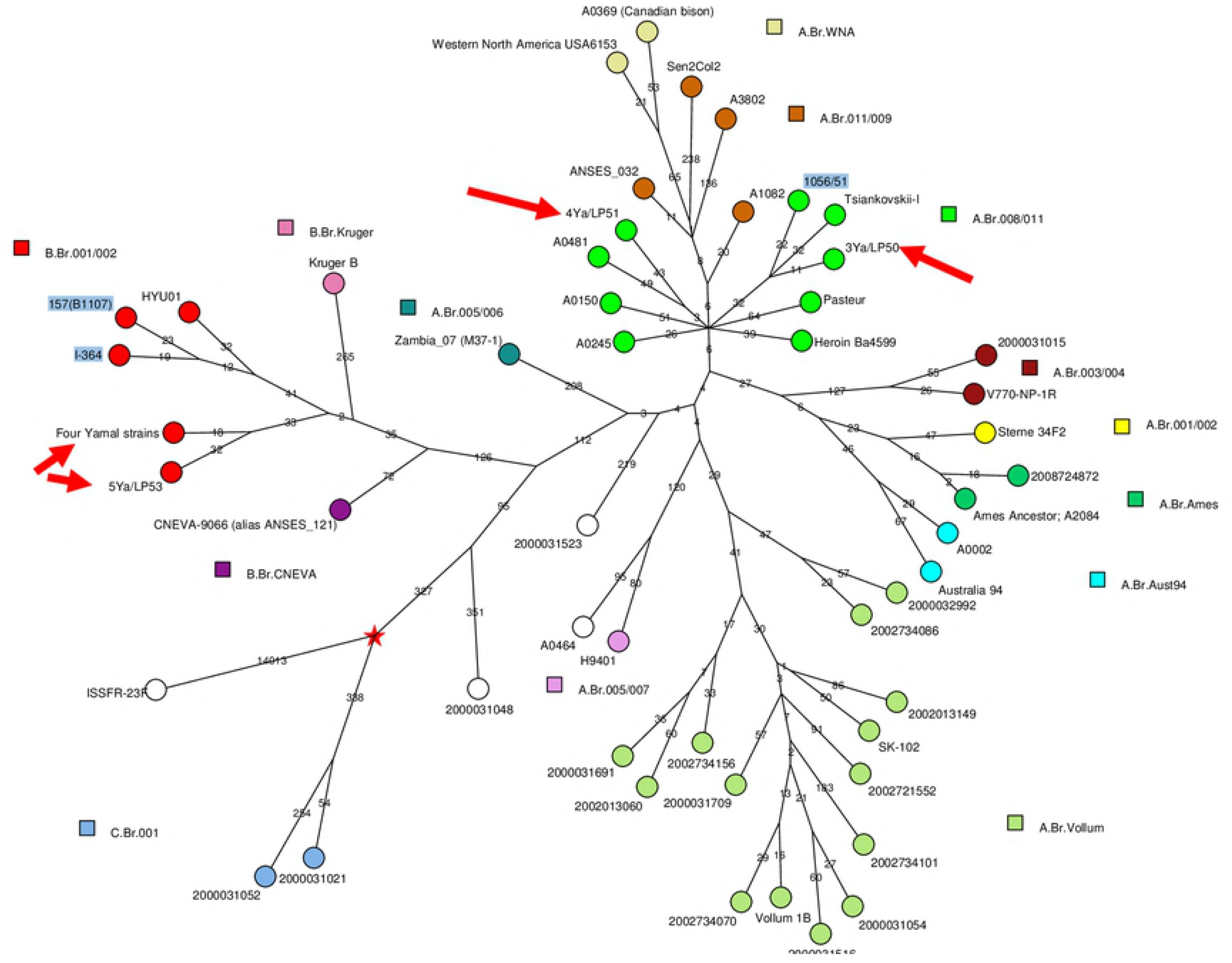
Position of the Yamal and Yakutia strains in the global *B. anthracis* phylogeny. Strains representing the main lineages including the different canSNP lineages were selected. Red star: the tree is rooted with *B. cereus* strain ISSFR-23F. Each circle is labelled with the corresponding strain name. The color code reflects the canSNP lineage. The very rare lineages defining the most ancient currently known splits have been found only in North America. The Yamal and Yakutia strains are arrowed. The number of SNPs constituting each branch is indicated. A logarithmic scaling was used in order to visualize the shorter branches.

Figure 2 is a close-up on the group including all available genome sequences assigned to B.Br.001/002 and sub-lineage B.Br.Kruger. B.Br.001/002 is split in two parts, one part including the B.Br.Kruger sublineage predominant in South Africa and the other part including strains from Northern Russia, Estonia and Korea. The “Kruger” group is characterized by relatively long branches. From the split to the tips of the tree, the “Kruger” group expansion ranges from 256 to 368 SNPs. In contrast, in the same timespan the “Eurasia” group expanded by 60 up to 86 SNPs, i.e. a ratio of 4.25 between the two groups. This observation may suggest that the “Kruger” clade is the result of a secondary introduction to South Africa. The Yakut Ya5 and Yamal strains belong to the “Eurasia” group and are separated by 54 SNPs.

**Fig 2.**
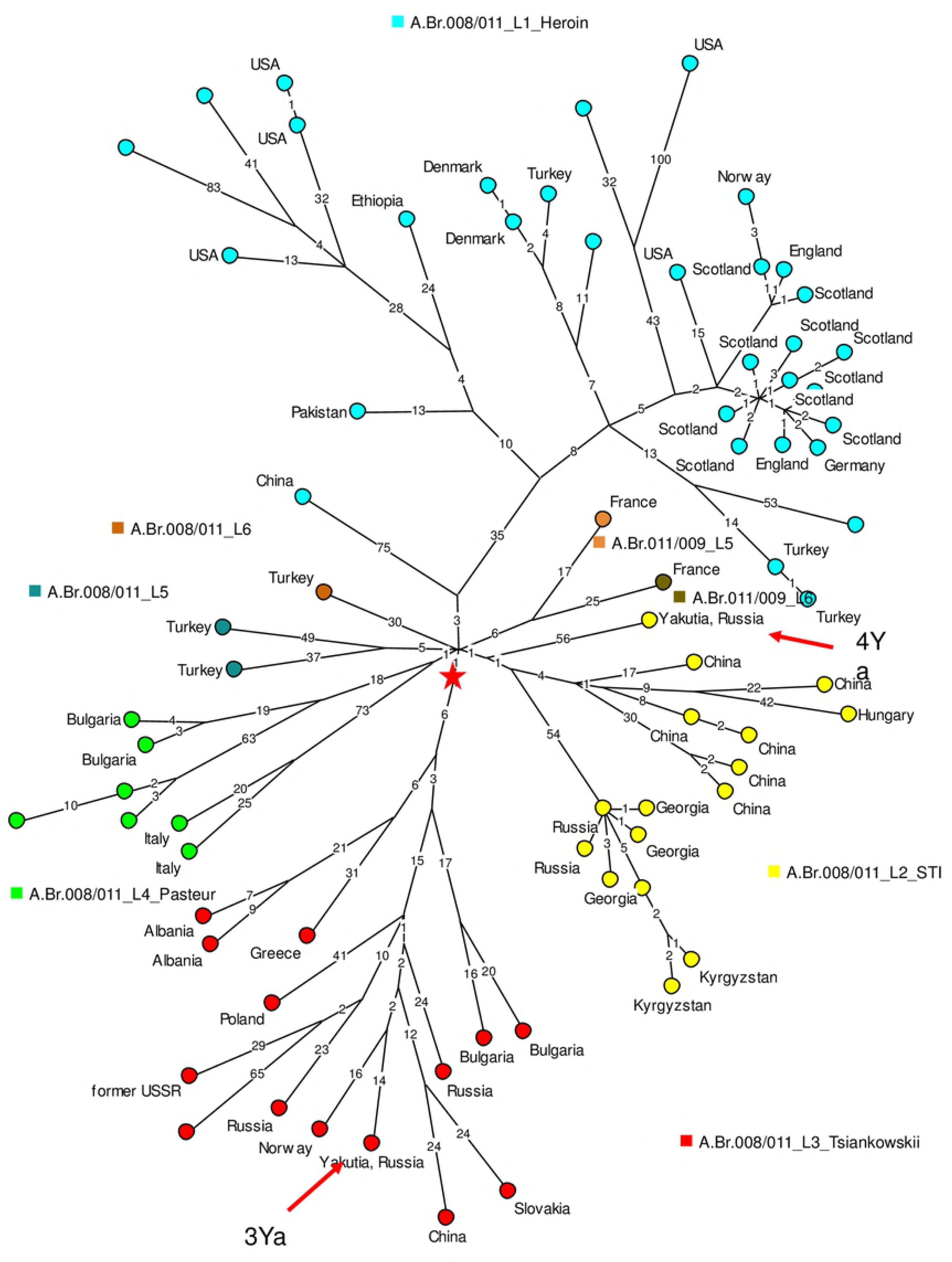
Focus on the B.Br001/002 and B.Br.Kruger lineages. The Ames ancestor reference genome is used to root the minimum spanning tree. Branch length (number of SNPs) is indicated. Geographic origin of each strain is displayed when known.

The two other Yakut strains, Ya3 and Ya4, belong to the Transeurasian radiation TEA 008/011 [32]. TEA 008/011 is a remarkable polytomy currently including seven lineages. Figure 3 displays a minimum spanning tree based on the whole genome SNPs detected among the 75 strains assigned to TEA 008/011. The branches are named as previously proposed [32]. Distances from root to tips differ by a ratio of 11 among the different lineages. The shortest branch with 17 SNPs represented by a strain from China, is observed in lineage L2_STI. The longest, with 194 SNPs, is observed in L1_Heroin. The most represented lineages are L1_Heroin, L2_STI and L3_Tsiankowskii. The L1_Heroin lineage was previously investigated in detail [33–35]. L1_Heroin contains one early split, three SNPs from the core of the polytomy. One branch with 75 SNPs is present in China, whereas the other branch is more complex in terms of geographic origin as it was isolated in many countries. The shortest branch (57 SNPs from red-star root to tip in Figure 3) within this lineage is defined by a cluster of strains recovered in different European countries from human patients infected via drug usage. The longest branch is the outcome of a recent split along the shortest branch. After this split, the relative rate of expansion was 28. The difference in length is not the result of horizontal gene transfer events, as the associated SNPs do not show evidence for clustering [36]. The geographic location of the reservoir is uncertain. Afghanistan is a likely option if heroin contamination occurred as part of the drug production or initial packaging processes. Other candidate countries are Turkey and Pakistan represented by strains defined by short lineages, or additional neighbor countries not represented in current *B. anthracis* databases. The L2_STI lineage contains three deep-branching sublineages. The first is defined by the Yakut 4Ya strain, the second is present mostly in China, and the third corresponds to the STI vaccine strain from Russia. The Tsiankowskii vaccine strain, the Sverdlosk 1979 strain [34], and the Yakutia 3Ya strain which is closest to a strain from Norway belong to lineage L3_Tsiankowskii lineage containing a single deep-branching lineage widely spreading in Russia and Eastern Europe, including Greece, Albania, Bulgaria, Poland. Lineage L4_Pasteur contains two deep-branching sublineages, one found in Bulgaria and the other in Italy in addition to the Pasteur I vaccine strain. Lineages L5 and L6 each contain a single deep branching lineage, observed in Turkey. The last lineage leads to TEA Br011 corresponding to the A.Br.011/009 polytomy [37, 38] found in France. In summary, eleven deep-branching lineages are detected within the seven-branch TEA 008/011 polytomy, nine of which with a strong geographic assignment, to Turkey (2), China (2), Russia (2, including Northern Yakutia, Siberia), Italy, Bulgaria and France. The root (red star) defined by the branching point of the Ames ancestor reference strain is located on L3_Tsiankowskii at a distance of one SNP from the center of the polytomy connecting the six other branches. This might suggest that Europe is the geographic origin of the A.Br.008/011 polytomy. However this argument is weak and extensive sequencing of additional A.Br.008/011 strains will be needed to establish the geographic origin of the polytomy.

**Fig 3.**
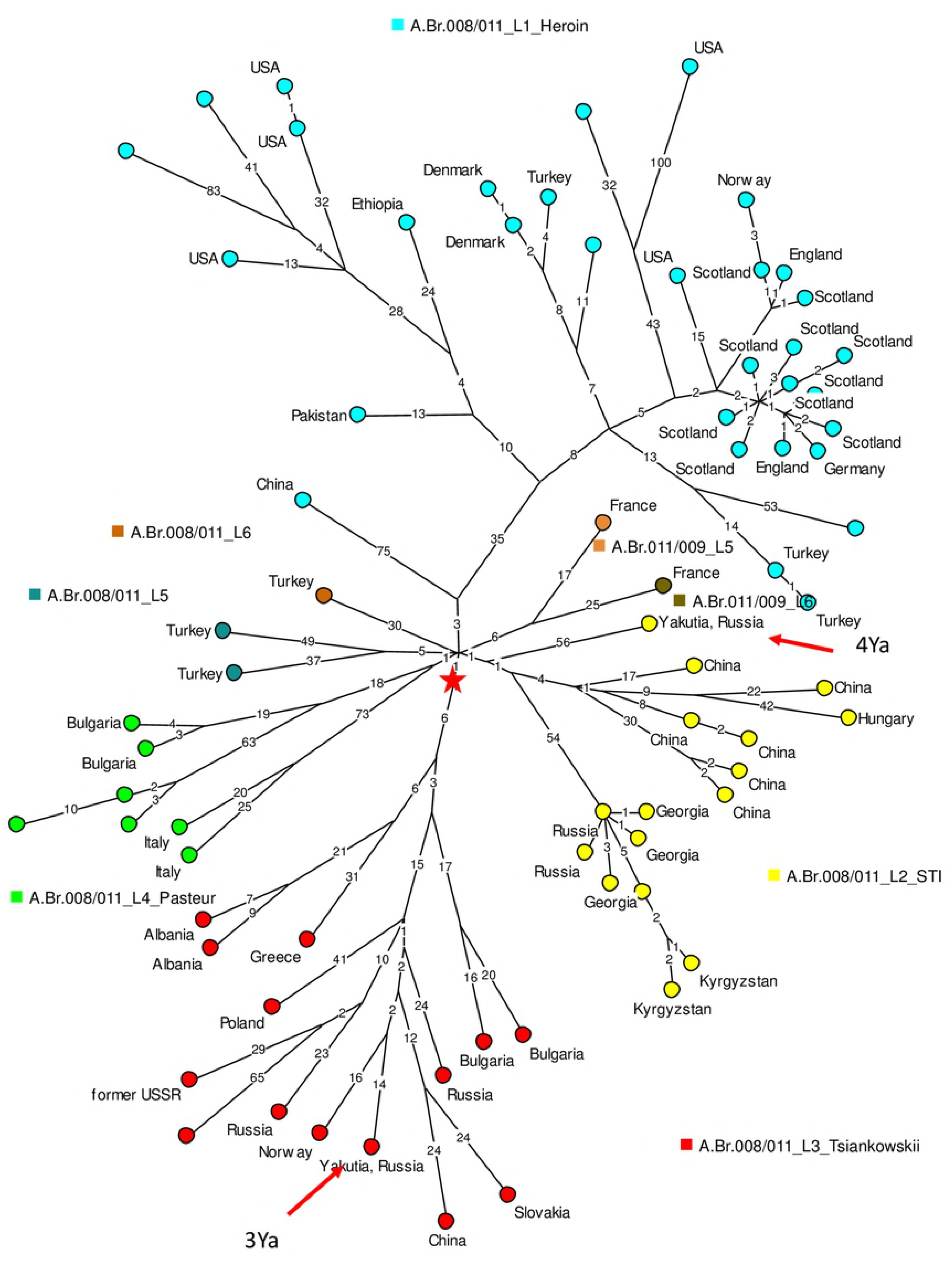
A.Br008/011 polytomy, minimum spanning tree, logarithmic scale. The color coding reflects the lineages within the A.Br.008/011 polytomy using the previously defined classification [32]. Geographic origin is indicated when known. The root (red star) is defined using the Ames ancestor reference genome. Two strains representing the A.Br011/009 lineage are included. The tree is based upon 1844 SNPs, and the level of homoplasia is 0.7% (the size of the tree is 1858).

Two representatives from A.Br.011/009 were included in Figure 3, in order to show the position of the root of the A.Br.011/009 polytomy located at a distance of six SNPs from the root of the A.Br.008/011 polytomy. Figure 4 shows the structure of the A.Br.011/009 polytomy, using the fifty-four read archives or genome assemblies assigned to A.Br.011/009. The polytomy comprises six branches numbered L1 to L6 in agreement with previous reports [37, 38]. The monophyletic A.Br.WNA lineage exclusively present and predominant in North-America emerged from A.Br.011/009_L2 and is represented by four strains in Figure 4 [28, 38–40]. Interestingly, the WNA lineage is branching from the Canadian strain A0303 which shows the ancestral state for the A.Br.WNA canSNP, i.e. still belongs to A.Br.011/009 [39]. This is in agreement with the report by Kenefic et al. demonstrating an introduction of WNA in the USA from Canada, and a progressive evolution from north to south. The West Africa clade from Guinea and Côte d’Ivoire [41] is branching out from lineage L2 at the same position as the WNA and Senegal-Gambia [42] clade. Apart from two Argentinian strains belonging to the L3_Pasteur II vaccine sublineage and one strain from the USA defining a recent split with a long branch within lineage L4, all the other A.Br.011/009 strains are from Italy and France. Italian strains define two deep lineages within lineages L1 and L2, which split at a distance of one SNP from the root of the A.Br.011/009 polytomy. After the split, the length of the expansion is very similar along the French and Italian sublineages (Fig 4). For instance the total length of the French L2 lineage is 26-33 SNPs, compared to 24-28 SNPs in the Italian L2 lineage.

**Fig 4.**
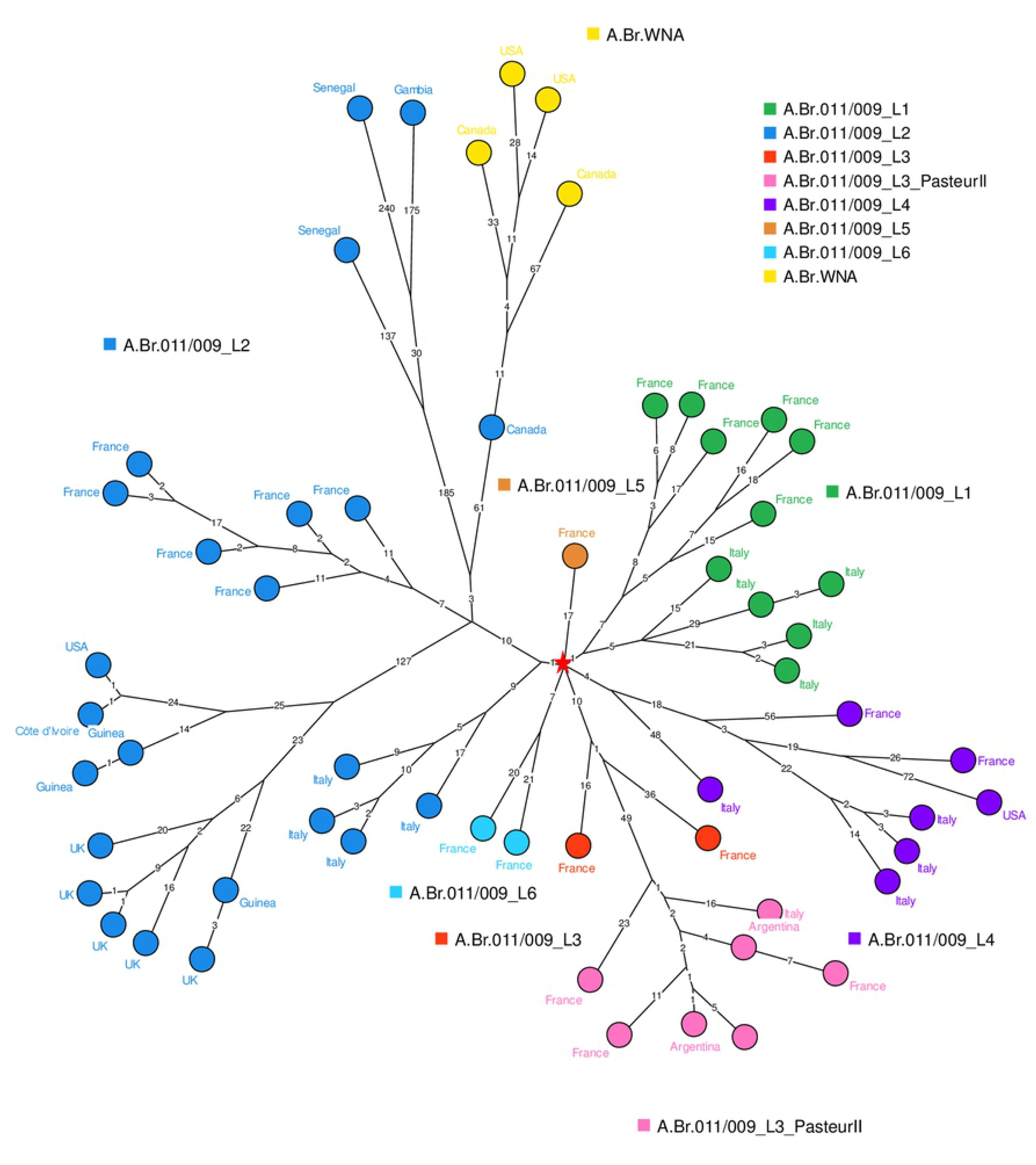
A.Br011/009 polytomy, minimum spanning tree, logarithmic scale. The color coding reflects the lineages within the A.Br.011/009 polytomy using the previously defined classification [38]. Within L3, strains derived from the Pasteur II vaccine are colored in pink. Geographic origin is indicated when known. The root (red star) is defined using the Ames ancestor reference genome.

Consequently, the analysis of currently available A.Br.011/009 sequencing data strengthens the previously reported view of the A.Br.011/009 polytomy, characterized by a limited geographic distribution of the “slowly evolving” lineages contrasting with the very fast expanding lineages observed in West Africa and North America [38]. After the North American-West African split shown in Figure 4, French strains from lineage L2 expanded for a further 15 to 22 SNPs when not including two strains obtained from the Collection de l’Institut Pasteur (CIP) which might have been extensively cultivated and define slightly longer branches. In contrast, the length observed towards Côte d’Ivoire-Guinea, Senegal-Gambia, and WNA are 167-177 SNPs, 323-458 SNPs and 112-142 respectively, i.e. ratios of 8 to 20. The currently available data provides evidence for two independent introductions of *B. anthracis* from the A.Br.011/009 polytomy in North America, one from the L2 lineage via Canada, and the second one from the L4 lineage. The second introduction is represented by a unique read archive SRR5811139 derived from a strain collected in New Hampshire. This second introduction occurred after L4 had already expanded for at least 2/3 of its current length whereas the first introduction from L2 occurred after L2 had expanded for 1/3 of its current length.

## Discussion

### The Yamal 2016 anthrax outbreak and implications

Anthrax is endemic in most of Russia, including Yamal. At the beginning of the Yamal colonization by the Russian Empire in the 17-18 centuries, cases of this disease were reported. The first outbreak was recorded in 1760. From 1898 to 1931, 66 epizootics were described, during which more than a million deer died. In the 1940s, the whole reindeer livestock was vaccinated. In subsequent years, the number of vaccinated animals was lower, for example, in the 1960s, from 65 to 82% of the total number of deer was vaccinated. This proved sufficient to prevent epidemics. Thus, a centuries-old cycle of anthrax circulation in the tundra of the Far North was interrupted [43].

The absence of outbreaks suggested that the soil in the tundra was sanitized and no longer contained anthrax spores. In 1968, 360 soil samples from places of recorded mass death of reindeers were examined and no *B. anthracis* strains could be recovered [44]. This suggested that tundra soil conditions (pH 3-5, humus content lower than 3%) are unfavorable to maintain the viability of the spores. In 2007, deer vaccinating was stopped. During June-July of 2016, the air temperature in the Yamal epidemic area in was 5-9 degrees higher than normal, and did not fall below 18 C. The soil reached a temperature of 25 C at a depth of 10 cm and 7 C at a depth of 1 meter. This was combined with a very small amount of rain precipitation [45]. Apparently such an anomalous warm climate led to the thawing of permafrost, and viable *B. anthracis* spores could be exposed to the surface [3, 46].

According to the testimony of reindeer herders in epidemic area near Pisyoto lake, herds from all focus of infection had been driven through the same place in the tundra. The thawing of permafrost provoked a landslide of a hill on the bank of the river. This could explain the finding of a single MLVA and wgSNP genotype. Unfortunately since all the helicopters in the region were being used to transport people and cargo, it was not possible to visit and sample the place (in this region roads are absent; movement is possible only by sledges with a team of reindeers or by helicopter). Along these lines, two migration ways of the pathogen might be proposed: washing out of bacterial cells from deep soil layers to the ground surface, or exposure of a deep soil layer to the surface due to thawing and landslide. At the same time, the reindeers were weakened by the heat, which could have increased susceptibility to infection. Observations in the focus of the disease and a survey of veterinarians and reindeer herders gave grounds to assume that infection could occur not only by spores but also by vegetative cells. In several cases, a sick reindeer could start recovering and stand on its legs after receiving a single dose of antibiotic, subsequently licked the muzzle of healthy animals, which fell ill and died within 24 hours.

Consequently, we cannot exclude a simultaneous spreading of infection from several isolated foci triggered by the exceptionally hot weather. A preceding large-scale epizootic, spread widely in the region and conserved in permafrost in multiple soil foci might explain the observed genetic homogeneity of the strains collected during the present outbreak. In our opinion, this alternative is not the most likely. Unfortunately, strains from previous outbreaks were not preserved in collections. Therefore, there is no way to compare the strain isolated in 2016 with strains previously circulating in the region, and, accordingly, to precisely estimate the length of time during which the pathogen spores were stored in the soil. Given that the last outbreak of anthrax in Yamal was registered in 1941, it is likely that the spores remained viable in the soil for at least 75 years.

Considering the scale of the epidemic and the costs of countering it, the time interval between the beginning of reindeer disease and the beginning of antiepidemic measures may seem too large. In the case of a faster response, the epidemic could be less severe. Several factors played a role in this situation:

- Presence of a large number of unvaccinated reindeer (Yamal reindeers population reaches 800 thousand), susceptible to infection and weakened by heat).

- Lack of experience in anthrax diagnosing - the last outbreak of anthrax was recorded in this region in 1941, and local veterinarians had no experience of this infection before 2016.

- Nomadic mode of cattle breeding - even a small herd can eat the limited tundra vegetation very quickly, thus breeders have to drive the herd to another place. Nomadic cattle breeding cover much larger territories than settled cattle breeding. The migration routes of different herds may cross each other. In the case of anthrax, this can rapidly increase the epidemic area, and contribute the infection to herds that migrate through the territories where sick animals were driven.

Also the establishment of the scale of the accident and the timely examination of the corpses was hampered by the fact that reindeer herders migrated and could not always bring dead deer for inspection. The delay in identifying patients with anthrax was also due to lack of awareness of the local population about the dangers of anthrax and about its symptoms. Local populations, even with symptoms of the cutaneous form of this disease, did not pay attention to them and considered themselves healthy. This situation arose from a set of reasons. The traditional way of life of reindeer breeders, transport and communication isolation of nomadic reindeer herders from cities is associated with a more limited access to medical care. The usual high prevalence of furunculosis prevented timely detection of anthrax affections on the skin.

All these factors favored the spreading of the infection before medical and veterinary organizations were alerted. During a hidden period, the infection may have been carried by transport (helicopters and ships), along with the goods and people who visited the outbreak, where the animals infection happened first. Unfortunately, it is not possible to retrace retrospectively the movement of people and transport between the foci of infection.

The occurrence of anthrax outbreaks after a 75-year break demonstrates that the decision to stop the vaccination was premature, in line with similar independent observations in Georgia [47]. Permafrost turned out capable to conserve viable microbial spores for a long period and figuratively speaking to be a reservoir of infection. Under favorable climatic conditions, these spores proved able to migrate to the surface of the soil and initiate new infection cycles.

If the Yamal strains were isolated during a large-scale epidemic, then the finding of the Yakut strains in a seemingly random place is very noteworthy. In the absence of historic record of anthrax outbreak in this area, there was no reason to expect the finding of *B. anthracis* spores in the soil. The microbiological investigation was prompted by the paleontological finding of cave lions leading to the serendipitous finding of *B. anthracis*. Three different genotypes from soil samples taken from a depth corresponding to the upper layers of permafrost down to four meters were characterised. One genotype was very close to the Yamal genotype and belongs to the B cluster of *B. anthracis* whereas the two others belong to the A.Br.011/008 polytomy.

### Tentatively dating the emergence of the A.Br.011/008 polytomy

The monophyletic, strictly clonal evolution of *B. anthracis* implies that the whole species derives from a unique progenitor [48]. Africa is sometimes proposed as being the “cradle” of *B. anthracis* [49]. The strongest argument in favor of such an origin may be the discovery in Africa of additional *B. cereus* lineages causing an anthrax-like disease [48, 50, 51]. However, this does not help dating or locating the origin of modern lineages, even when these lineages display a strong geographic preference. A tentative dating of the Most Recent Common Ancestor (MRCA) of the *B. anthracis* species to 13,000-27,0000 years was previously proposed based on average mutation rate and estimates of infection cycles per year [28]. The *B. anthracis* species may be much older than the MRCA defined by known lineages.

The dating of nodes along the *B. anthracis* phylogeny is particularly difficult and challenging because of the irregularity of its evolution due to its ecology [28, 32, 39]. In contrast with other pathogens such as *Mycobacterium tuberculosis* [36, 52, 53], branch length does not correlate with time, as investigated and discussed in detail by [32]. Rather branch length most likely reflects the number of infection opportunities per year [28] or more rarely a mutator phenotype [54]. This number is expected to increase when *B. anthracis* encounters a favorable ecological niche. Each split in the tree reflects the colonization of a new ecotype allowing the fixation of an additional, independent lineage. Usually such a split will result from geographic spreading, and the new ecotype may be the one with the longest, faster evolving branch, at least when a significant difference is observed, reflecting the arrival in a new, naive environment. In this context, polytomies constitute exceptional opportunities to try and propose dating points. Polytomies result from the sudden colonization of multiple new ecotypes which may reflect exceptional environmental changes. Such changes may have an anthropic origin and it may be easier to associate a polytomy with major historical events. Two large polytomies have been described so far in *B. anthracis* the A.Br.008/011 [32] and the derived A.Br.011/009 polytomies [37]. They constitute the “TransEurAsia” (TEA) subclade [32].

The branching of fast-evolving lineages from the same position within a unique sublineage of the A.Br.011/009 polytomy towards both West Africa and North America indicated that the contamination was exported from a geographically limited region having exchanges at the same time with both continents (Canada, Senegal-Gambia). France, seventeenth century was proposed as the most likely spatiotemporal candidate [38]. Based on the proposed dating, a tentative dating of the most recent ancestor of the A.Br.011/009 polytomy to the One Hundred Years War between France and England, AD 1350-1450 was further hypothesized [38].

However, this proposition does not fit well with the present report showing a remarkable intermingling of Italian and French strains early during the emergence of the A.Br.011/009 polytomy. Rather this intermingling would indicate that the A.Br.011/009 emerged in a time of conflicts between Italy and France, rather than France and England, before the seventeenth century. Years 1250-1300 and years 1450-1550 are two candidate periods. Battles at that time involved thousands of horses. We speculate that such events provided major opportunities for contaminations of both cavalries with the same population of strains. These large “flocks” will then spread the contamination on their way back.

In contrast to the A.Br.011/009 polytomy, the A.Br.008/011 polytomy is characterized by a remarkable geographic spreading. Rare, deep branching lineages are observed in Europe (Italy, Bulgaria, France) as well as Turkey, China and Yakutia (the permafrost strains, this report). There is one major event in human history prior to the sixteenth century which could explain such a distribution, the Mongol conquests during the thirteenth century [55, 56]. Chinghis Khan assembled an empire extending from Northern China to the East side of the Caspian sea. The death of Chingis-khan in 1227 triggered the gathering of the Chingizids Armies for the election of Ögedei as new khan, and the death of Ögedei in 1241 eventually triggered a new gathering in 1246. The conquests, powered by horses, involved long-distance displacement of tens of thousands of horses. The immediate successors of Chingis-khan invaded Europe as far as Hungaria in 1237-1242 and Anatolia (Turkey) in 1241-1243.

The Mongol Empire began to disintegrate in the second half of the 13th century. But despite this, the territory from China to Eastern Europe remained under the rule of Chingizids, and Eastern Europe was constantly subjected to raids from the Golden Horde for the purpose of plundering, or simply the presence of military contingents participating in wars. Thus, a common political space was established, ensuring relatively large movements of people and animals between Asia and Europe, which could further contribute to the relatively rapid and unhindered transfer of pathogens of infectious diseases between these regions. In addition to military operations, the Mongols organized a Yam - chain of relay stations at certain distances to each other, allowing to replace horses for messengers or messengers themselves, and making possible to deliver cargo and documents within weeks over long distances. This postal system also could promote rapid spread of infections, but to a much lesser extent than the massive movements of armies driven by horses.

The presence of Chingizids in the European region ceased when Russian Tsar Ivan the fourth (Ivan the Terrible) conquered Western states, which were formed after the split of the Golden Horde - the Kazan Khanate in 1552, the Astrakhan Khanate in 1556 and the Siberian Khanate in 1582-1598. After that, the only post-Mongolian state remained the Crimean Khanate, which regularly carried out raids on Russia and Poland (the territories of modern Russia, Ukraine, Belarus, Lithuania, Latvia, Estonia and Moldova) until it was conquered by Catherine the Great in 1783. Despite the active raid policy of the Crimean Khanate and the constant use of the Tatar contingents by the Moscow tsars in the course of constant wars in eastern and northern Europe, the continuity of migration routes of nomads from Asia to Europe was broken.

Consequently we propose here that the root of the A.Br.008/011 polytomy corresponds to a *B. anthracis* ecotype present in the Mongolian armies between the first half of the 13^th^ century and the middle of the 16^th^ century. We particularly favor the first half of the 13^th^ century as the period when *B. anthracis* could have been transported in a short time-frame by the animals associated with the Mongolian armies, particularly war and led horses in all geographic areas covered by the deep branching lineages of the A.Br.008/011 polytomy, i.e. from China to Hungaria. After that time, we speculate that the split of the Mongolian empire would have significantly hindered the spread of the infection.

In contrast with this relatively precise dating hypothesis, we see at present no clue regarding the geographic origin of the ancestor of the A.Br.008/011 polytomy. The contamination of the Mongolian army might have occurred in many locations, including Eastern Europe.

### Apparent contradiction between the dating of the emergence of the A.Br.011/008 polytomy and of the permafrost samples

Under the proposed hypothesis, the A.Br.011/008 polytomy can be conservatively dated from the early 13^th^ century to the middle of the 16th century. The deposition of the A.Br.011/008 strains including the one represented by 4Ya recovered from permafrost at a depth of two and three meters would be posterior to this period. The 3Ya sublineage is even more recent, the relative length of the branch is less than half the total length of lineage L6_Tsiantowskii to which it belongs.

The Yakoutia permafrost soil samples were taken from alluvial (river) sediments - there is a wide flat valley with bayou lakes. Probably this corresponds to holocene sediments (age about 10,000 years), frozen as they accumulated. A layer of permafrost formed simultaneously in Yakutia and Yamal 20-40 thousand years ago. Radiocarbon analysis of other sediments sampled in Uyandina riverside and other places of Abyisk district at the same depth, indicated that they were 3-10 thousand years old [57]. On average, in Yakutia, the depth of seasonal thawing does not exceed 2 - 2.5 meters. Therefore we expected that strains of *B. anthracis*, isolated from the permafrost, would be significantly older. However, the accidental nature of the exceptional paleontological finding in Yakutia, and the extraction of paleontological material and soil samples by prospectors rather than professional paleontologists or geologists, may be responsible for a weak geological information about the soil layers from which the strains were extracted. The finding of *B. anthracis* was quite unexpected and was triggered by the cave lions investigation, and some time passed between the excavation of the cave lions and the soil sampling.

We also carefully examined the extent to which the present findings could be the result, or be affected, by contaminations of different kinds. However, the possibility that the strains were introduced into the soil samples under study as a result of contamination during work is most unlikely. Drift of spores from the surface is unlikely, because of the absence of reported cases at the sampling site recently and in view of the absence of *B. anthracis* spores in the upper (minus 1 meter) sample. Most importantly the initial identification of *B. anthracis* was made in a laboratory which does not work with pathogenic microorganisms and does not maintain such strain collections. The contamination in SRCAMB is also very unlikely as it would require a simultaneous contamination with three different strains in only three samples. In addition and most convincingly, whole genome sequence analysis of strains from the SRCAMB collection showing a similar MLVA genotype demonstrated that these are definitely different strains.

A number of conclusions can thus be robustly drawn. We looked for the presence of *B. anthracis* spores over a depth of six meters, going from one meter below surface down to the cave lion kittens. The state of preservation of the kittens indicated that the bodies, and the corresponding permafrost layer, remained frozen for thousands of years. Under these conditions, the most parsimonious explanation for the finding of *B. anthracis* strains only in the upper layers is that *B. anthracis* was not present in the region at the time of the death of the kittens, 5000 to 10,000 thousand years ago. Rather *B. anthracis* arrived relatively recently in three occasions represented by three distinct lineages. This diversity cannot be the result of pathogen evolution in the soil. The three genotypes are clearly positioned in different places of the *B. anthracis* evolutionary tree. Thus, all three strains most likely hit the ground and were conserved at different times, during various epizootics. The lack of a clear stratigraphy, that is, the isolation of isolates that represent a single genotype from soil samples from different depths, and more importantly the finding of spores lower than expected from the proposed dating would imply that microorganisms are able to migrate in permafrost. This would be compatible with current knowledge on permafrost [58, 59].

### Explaining the close relationship between the Yamal and Yakoutia *B. anthracis* strains

One surprising observation in our study is the fairly close genetic relationship of the Yamal isolates with the Yakut strain 5Ya. Between the regions where these strains were isolated lies a distance of about 2 thousand kilometers. However, despite their remoteness from each other, they are located at similar longitude, and the ecosystems located in them are almost identical. In this connection it can be assumed that these strains are representatives of a certain "tundra" population of *B. anthracis*, spread on a significant territory of Northern Eurasia, at least in the past. In this case, the circulation of strains could be ensured by migration of ungulate populations, primarily reindeer.

The territory of Yakutia was inhabited by modern humans at least from the Mesolithic, but before the beginning of the 2nd millennium AD it was inhabited exclusively by tribes of hunters and fisher-men. The first population of herders, the ancestors of modern Yakuts, migrated here only at the beginning of the second millennium from the Baikal region. They were livestock breeders, bred cows and horses (in the Baikal region they also bred sheep and camels, but it is impossible to breed them in the territory of Yakutia due to the severe climate).

The conquest by the Mongols of southern Siberia, slightly increased the migration movement from Baikal region. Tribes with domestic reindeer that came from the south of Siberia (Baikal region), by the middle of the second millennium AD reached the territory of western Siberia, which includes the Yamal Peninsula, and the north of Eastern Siberia - Yakutia. In western Siberia, these were the ancestors of the Nenets, in the eastern of the Evenks. Theoretically, it can be assumed that in the middle of the 2nd millennium AD those and others could contact somewhere in the area of the Yenisei basin, south of Taimyr, for example, in the Turukhansk district, from where a route to the Yamal is possible. At that time reindeers were mainly used for transportation and the number of domestic deers was low. Until the 17th century there were no large herds, the maximum livestock in one farm could reach one hundred heads of deer. Large-herd reindeer herding developed only after the colonization of Siberia by Russians, beginning from the 17th and 18th centuries. Currently, the number of herds in one farm reaches several thousand heads. More knowledge on *B. anthracis* strains present in Northern Europe and Siberia will help better understand the history of *B. anthracis* spreading among reindeers.

Only in the middle of the 20th century, in connection with the beginning of mass vaccination of reindeer and the introduction of veterinary and sanitary control, obstacles arose for the free distribution of *B. anthracis* in the tundra zone. However, considering the ability of *B. anthracis* to form endospores and the presence of permafrost, capable of preserving these spores, further increasing the period of their viability, multiple soil foci could have formed by this time scattered over a vast territory. This territory is very little involved in economic activity and is extremely poorly populated (for example, the average population density in Yakutia is 0.3 people per square kilometer; since two thirds of the population lives in cities, the density in rural areas is only about 0.1 person per square kilometer). This makes not only sanitation, but even detection and recording of such foci an impossible task. The events of the summer of 2016 in Yamal have shown that such foci retain their epidemiological potential for a long time, and under favorable conditions, primarily in the thawing of permafrost due to local or global warming, they can become a source of infection, causing large-scale epidemics, resulting in casualties and significant economic costs. A reactivated focus may remain active for some years. In 2017 32 samples from the Yamal 2016 epidemic area were tested. Two samples - ash from the place of a dead deer burning (lat. N68.24010, lon. E71.01.200) and the biological material from not completely burnt deer (lat. N68.24989, lon. E70.44435) – containing PCR detectable *B. anthracis* DNA detected in PCR. Virulent bacteria could be cultivated from both samples. The MLVA genotype was identical to the strains previously isolated in the epidemic area in 2016 [60].

If spores were able to keep viability during a year on the soil surface, then there is little doubt that during seasonal snow melting they could be spread over a large area and penetrate into the depths of the soil. Thus, in the tundra areas, where at least once an anthrax outbreak was recorded, it is necessary to conduct continuously anti-epidemic measures, such as vaccination of livestock and herdsmen, as well as maintain the readiness of medical and veterinary institutions for the diagnosis of anthrax and emergency measures for detecting the disease.

## Conclusion

In summary, we have detected three independent events of *B. anthracis* introduction in Northern Yakutia, stored at different depth in the permafrost. In the proposed model, the third and most recent introduction, detected at minus 2 meters, occurred as a side effect of Russian conquests and development of agriculture in the 17^th^-18^th^ century. The second introduction detected at minus 2 and minus 3 meters, would be the byproduct of Yakut’s population migration from Lake Baikal area in the 14^th^-15^th^ century. The first introduction detected at minus 4 meters, cannot be dated precisely but the location in the permafrost may indicate that it is not more than a few centuries older than the second introduction. We propose to date the emergence of the A.Br.008/011 polytomy to the first half of the thirteenth century, in relation with the Mongolian conquests.

## Acknowledgements

We wish to express our gratitude to Vitalii Kharkov, the Chief doctor of Center for Hygiene and Epidemiology in the Yamalo-Nenets Autonomous District for his help in organizing of sampling, Dr. Alla Ryazanova from Stavropol Anti-Plague Institute for her role in the clinical and laboratory diagnosis of anthrax during the Yamal anthrax epidemic and Dr. Oktyabrina Sofronova, the head of the Laboratory of especially dangerous infections of Center for Hygiene and Epidemiology in the Republic of Sakha (Yakutia) for samples from paleontological discovery point sent to SRCAMB. Also we show our appreciation to Dr. Victor Karlov from Department of Ethnology, Faculty of History, Moscow State University for kindly provided historical advice and to Dr. Anatoli Brouchkov, the Head of Geocryology Department, Faculty of Geology, Lomonosov Moscow State University, for his geology advice.

## Supporting information caption

**S1 Table. The MLVA profiles of described strains**

**S2 Table LD50 values of studied strains for mice and guinea pigs**

